# Rapid Evolution of Colistin Resistance in a Bioreactor Model of Infection of *Klebsiella pneumoniae*

**DOI:** 10.1101/2022.10.03.510619

**Authors:** Juan-Carlos Jiménez-Castellanos, Bartlomiej Waclaw, Alison Meynert, Thamarai Schneiders

## Abstract

**Objectives:** We seek to understand the dynamics of *de novo* resistant mutations arising during colistin treatment of an antibiotic-naïve population of *K. pneumoniae*.

**Methods:** We used a bioreactor model of infection and colistin treatment against the colistin susceptible *K. pneumoniae* Ecl8. Whole-genome sequencing and MIC profiling was used to characterize genetic and phenotypic state of the bacterial culture at three time points: before treatment, immediately after regrowth following challenge, and at the end point of the experiment. A mathematical model based on the birth-death process was used to gain further insights on the population dynamics of emerging resistant variants.

**Results:** We find that, after an initial decline, the population recovers within 24h due to the evolution of highly resistant clones which exhibit MICs >100-fold higher than the parental strain. Recovery is caused by a small number of “founder cells” which have single point mutations mainly in the regulatory genes encoding *crrB* and *pmrB*. The mutants arise through spontaneous mutations prior to colistin treatment.

**Conclusions:** Development of colistin resistance during treatment of *K. pneumoniae* infections is readily achieved if colistin is used as a monotherapy.

## Introduction

*Klebsiella pneumoniae* is an opportunistic highly antibiotic resistant Gram-negative pathogen, currently recognised by CDC and WHO as a severe threat to both human and animal health^1,2,3^. The emergence of pan-drug resistant variants of *K. pneumoniae* significantly limit treatment options, demanding an increased reliance on last-resort agents such as tigecycline, carbapenems and colistin^4,5^.

Colistin is a cationic polypeptide that electrostatically interacts with the anionic bacterial outer membrane, displacing calcium and magnesium ions from the lipopolysaccharide (LPS). In *K. pneumoniae*, where colistin resistance arises upon LPS modifications through the activation of the *pmrHFIJKLM* operon (also known as *arnBCADTEF*) resulting in the alteration of the LPS charge via the addition of 4-amino-4-deoxy-L-arabinose to lipid A^6,7^. The activation of *pmrHFIJKLM* operon has been linked to multiple mechanisms which include loss-of-function mutations in the small lipoprotein MgrB and single point mutations within histidine-kinases (i.e., PhoQ, PmrB and CrrB) ^1,6^ leading to constitutive phosphorylation of the cognate response regulators (i.e., PhoP, PmrA and CrrA)^8–10^ and activation of the *pmrHFIJKLM* operon. Furthermore, expression of the PhoPQ, PmrAB and CrrAB systems can also result in the production of connector proteins such as PmrD or CrrC, further amplifying the activation of *pmrHFIJKLM* operon ^8^.

Given the multi-mechanistic basis of colistin resistance in *K. pneumoniae*, we sought to understand which *de novo* chromosomal mutation(s) would arise under typical treatment scenarios^11^. Using a continuous culture bioreactor to simulate colistin treatment of a *K. pneumoniae* infection^12^ in combination with whole-genome sequencing, phenotype characterisation, and mathematical modelling enabled us to gain insights into the process of *de novo* resistance emergence and selection of resistant mutations, which would be difficult to study in human or animal models.

We show that *K. pneumoniae* develops genetic resistance to colistin within 24h at a clinically relevant concentration of the antibiotic (10mg/L) based on standard pharmacokinetic and pharmacodynamics parameters use for colistin regimens^11,13–15^. Our results show that resistance emerges mostly through mutations within CrrAB and the PmrAB locus but not in MgrB. Over time, we observed some mutations being replaced by better-adapted variants. Our mathematical approach, which models the population dynamics of resistant mutants, reproduces the experimental data and provides insights into the experimentally inaccessible early stages of mutant emergence and growth. We find that colistin selects for the fastest-growing mutants from a large pool of resistant variants which exist at low numbers even before colistin exposure. Further application of our mathematical model shows that colistin resistance is likely to evolve at different bacterial loads and growth rates relevant for real diseases. Together, our approach suggests that intrinsic *de novo* mutations conferring clinically-relevant colistin resistance arise easily in patients during treatment, and underscores the importance of using colistin as part of combination therapy.

## Materials and Methods

### Bioreactor as a model of *K. pneumoniae* infection

We cultured *K. pneumoniae* Ecl8^16,17^ in an automated bioreactor, similar in construction to the turbidostat^18^ and the morbidostat^19,20^. The system consists of four identical bioreactors. In each bioreactor, bacteria grew in a cylindrical glass bottle (approx. 25mm internal diameter, approx. 40 mL max. volume) equipped with a magnetic rod driven from underneath by a multi-position stirrer (SciQuip GyroStir 10), a set of inlet and outlet tubes, and an optical sensor for turbidity measurements (Fig. 1). A set of miniature peristaltic pumps connected to media reservoirs delivered the media (LB and 50mg/L colistin in LB). A 4-channel peristaltic pump pumped out spent medium and the surplus bacterial culture. The outlet tube was positioned so that, with the outlet pump running, any surplus volume over the 25mL culture volume would be removed. Further details are provided in the supplementary information.

**Fig. 1.**
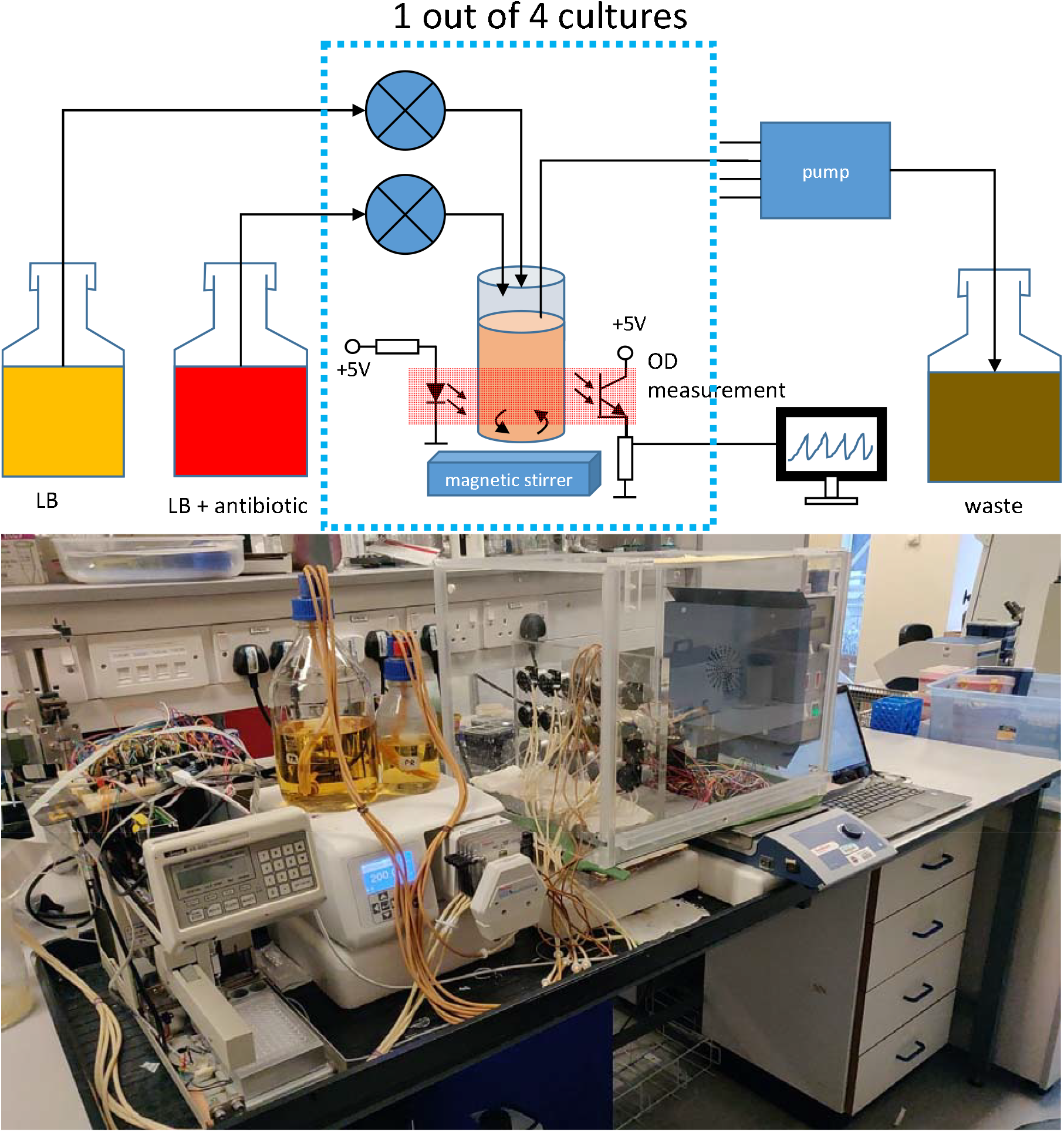
Automated culture system. Top: schematic diagram. The dotted blue box represents one of four bioreactors; all kept inside an incubator set to 37°C. Media and waste bottles are outside the incubator. Bottom: a photograph of the actual system.

An Arduino micro-controller connected to a PC, running a custom-written (C++) software to control the experiment, recorded the OD (every 2.5s) and the temperature (every 25s). The software initiated dilution with fresh medium every 1h or when the optical density exceeded OD_600_=0.5, or whichever happened first. To dilute, an appropriate volume of fresh media (LB only, or LB + colistin) was first added. Next, the outlet pump ran for 23s, long enough to pump out any surplus culture and reduce the total volume to 25mL. The volume of the injected fresh media was such that a fraction *f* = 0.66 of the bacterial culture remained in the system (or, conversely, a fraction 1 − *f* = 0.34 was removed).

An automated sampler connected to culture bottle 1 retrieved two 200 µL samples of bacterial culture every hour, and injected them into a 96 well plate with empty wells (sample 1), and wells with 100 µL 70% ethanol (sample 2), all covered with transparent seal to prevent evaporation.

### Evolution of resistance to colistin

We performed 8 experimental runs, E1-E8, with 4 independent cultures in each (total 32 biological replicates E1.1-E8.4). After inoculation with 200 µL of dense overnight culture, cells were incubated in the bioreactor without the antibiotic until the OD_600_ reached 0.5 for the first time, and for one hour afterwards. During the next dilution cycle, an appropriate volume of 50 mg/L colistin was injected to obtain the desired concentration (10 mg/L) in the culture. This bacterial concentration was maintained for the rest of the experiment by adding appropriate volumes of LB and colistin. The experiment was stopped between 24 and 26 h (different per run) post antibiotic challenge. We initially performed two pilot runs (P1.1, P2.1) to optimize the experimental protocol. The main difference of these runs to E1-E8 was the increased sensitivity of the optical density detectors, which caused saturation as the OD approached 0.5 resulting in lower OD values being reported than the true OD.

### Whole-genome sequencing

We performed WGS for 8 experimental runs **E1.1, E2.1**,**E3.1**,**E4.1**,**E5.1**,**E6.1**,**E7.1**,**E8.1**, at two time points: a cycle in which OD increased above zero for the first time during the regrowth phase, and a cycle close to the end of the experiment. In addition, we sequenced samples from the two pilot experiments (**P1.1, P2.1**) at three time points: the last pre-colistin cycle, early regrowth following challenge, and near the end. We also sequenced an overnight culture (OVC) grown in LB without colistin to use as a reference genome.

Genomic DNA was extracted using Lucigen Kit as per manufacturer’s instructions (Bioresearch, Lucigen, UK) and samples were sequenced by Novagene (Cambridge, UK) using an Illumina Novaseq platform with 150bp paired end reads generating 1GB of raw data (mean coverage 263x). Samples were aligned to *K. pneumoniae* Ecl8 reference genome^17^. Allele variants were called and their frequencies were obtained as described in SI Section “WGS/Data Analysis”. Table S1 lists all amino-acid changing variants and their frequencies.

### Minimum Inhibitory Concentration (MIC)

End-point samples from experiments **P1.1, P2.1** and **E1.1-E8.1** were plated onto colistin agar plates (10 mg/L). From each plate, three different colonies were picked to establish colistin susceptibility by microbroth dilution method according to CLSI guidelines^21^. Briefly, MIC experiments were carried out in 96-well flat bottomed microtitre plates (Sarstedt, UK) with LB plus colistin (concentration range 0.25 to 256 mg/L). 20 µL of bacterial suspension (OD_600_=0.01) was added to each well. Plates were incubated overnight at 37⍰°C for up to 20⍰h and absorbance (OD_600_) was read using a Fluostar Optima (BMG Labtech) plate reader.

Mathematical model. We used a model based on the birth-death process with mutations to simulate the population of bacteria in the bioreactor. At time *t* = 0 only colistin-sensitive cells were present. Sensitive cells replicated with rate *r*_*S*_(*t*)(1 − *µ*), and mutated into resistant cells with rate *r*_*S*_(*t*)*µ*, where *µ* ≪1 represented the per-cell per-generation probability of a colistin-resistant mutation. In the absence of colistin, resistant cells also grew with rate *r*_*S*_(*t*). In the presence of colistin, sensitive cells did not grow, but resistant cells replicated with time-independent rate *r*_*R*_. Both sensitive and resistant populations were periodically diluted (as in the experiment) by removing a fraction 1 − *f* = 0.34 of all cells.

*r*_*S*_(t) was time-dependent to account for changes in the growth rate necessary to reproduce the observed OD vs time curves from before colistin exposure (Fig. 6). Specifically, *r*_*S*_(*t*) was a piecewise function, each interval representing a period between two consecutive dilutions. To obtain it, we fitted the function *C* + exp(*A* + *Bt*) to the *OD*(*t*) in each dilution interval, and calculated the number of sensitive cells *N*_*S*_ = *VX*(C+exp(*A* + *Bt*)). Here *V* = 25ml and *X*= 4 × 10^8^ ml^−1^ is the proportionality factor representing the density of cells at OD=1, extrapolated from CFU counts of suspensions of *K. pneumoniae* of OD=0.1 and OD=0.5. We then determined the growth rate as *r*_*S*_(*t*) =*d* (ln *N*_*S*_(*t*))/*dt* = *B* (exp[*A*+ *Bt*])/(*C* + exp[*A* + *Bt*]).

*r*_*R*_ was assumed to be time-independent since we only simulated the model until the population recovered; OD was low in this phase of the experiments, and resistant bacteria were expected to grow exponentially with constant rate. *r*_*R*_ was determined as the maximum growth rate of the population following colistin exposure. In all simulations we used *r*_*R*_ = 1.7 *h*^−1^ which was the average regrowth rate in E1-E8 (Fig. S1).

The probability of mutation, *µ*, was obtained by finding *µ* such that the similarity (measured as the p-value of the Kolmogorov-Smirnoff test) between the experimental and simulated distribution of regrowth times was maximized.

## Results

### Colistin selects for highly-resistant mutants in a single-step

Upon exposure to colistin, cell density sharply decreased below the detection threshold in all cultures within 7 hours (Fig. 2). Nevertheless, all cultures eventually recovered pre-colistin cell densities 12-17 hours post-exposure. The time from exposure to regrowth showed little variation (Fig. S2, mean = 18.3 h, std. dev. = 0.9 h). Cells from the end-point of the experiment were highly resistant to colistin, with MICs of at least 64 mg/L and over 256 mg/L for some cultures (Table 1). Such a significant increase in resistance has been previously reported for *P. aeruginosa* exposed to colistin in the morbidostat, in which however, it took 10-20 days to evolve^22^.

**Table 1.**
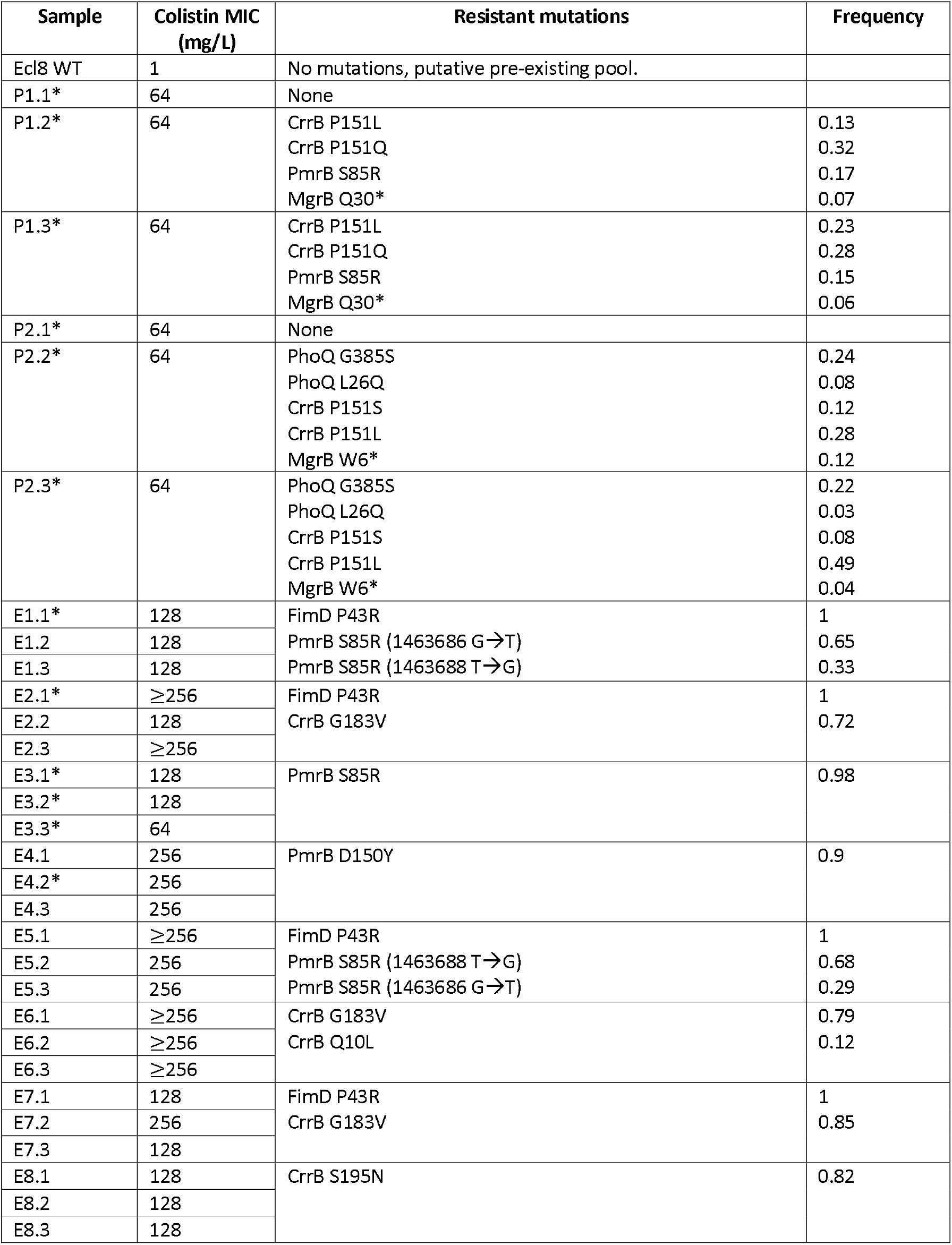
MICs of end-point samples, and mutations detected in these samples, gene name, frequency. Asterisk “*” denotes “dry” colonies (large, flat colonies).

**Fig. 2.**
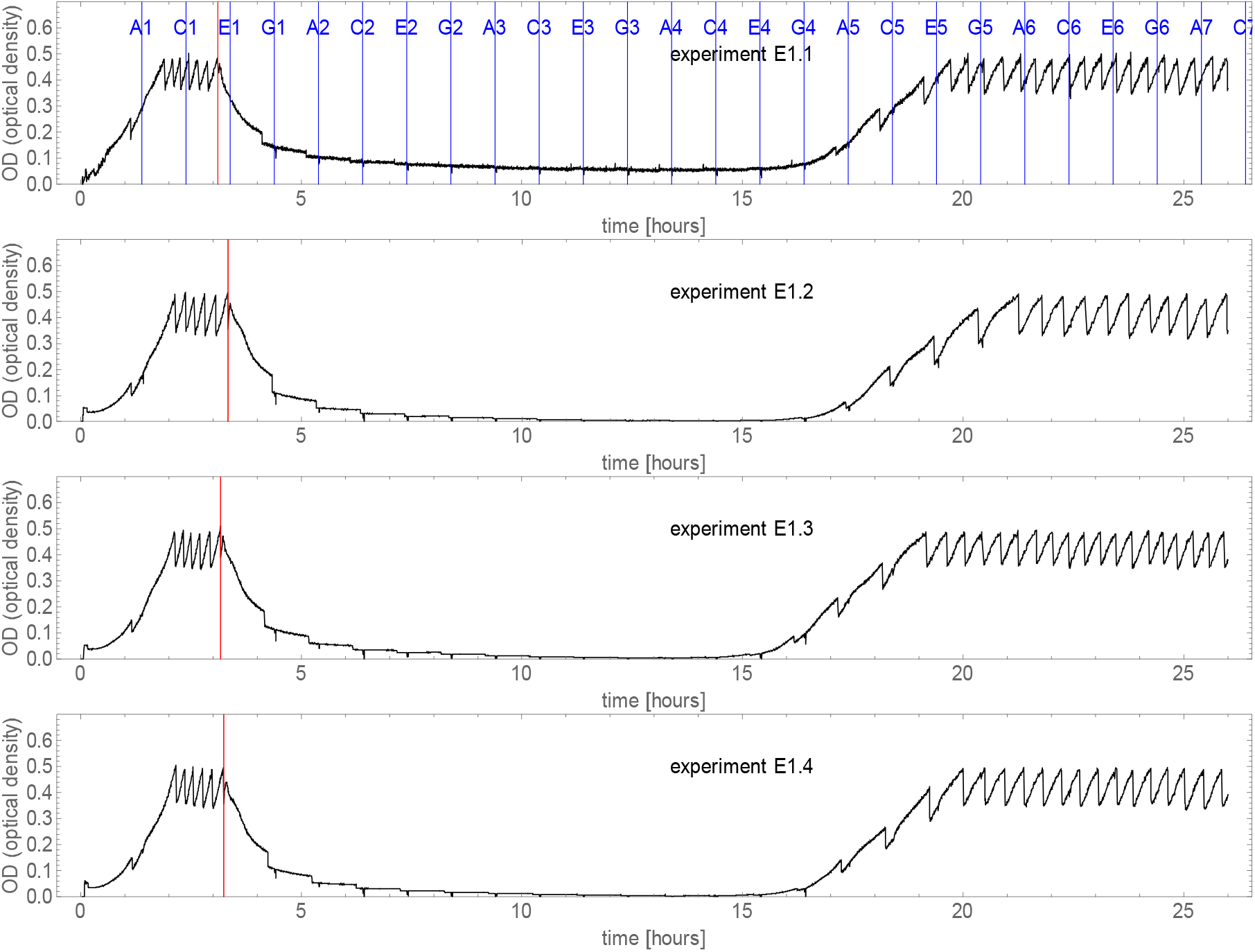
Representative examples of OD vs time curves for four experiments (run E1). Red line indicates the beginning of colistin challenge. Blue lines show the times at which the culture was taken by the autosampler, with sample labels denoted A1, C1, E1, We performed WGS for samples C5 and E6 in experiment E1.1.

The observed behaviour is not specific to Ecl8; Fig. S3 shows the same consistent regrowth for *K. pneumoniae* 52145^23^. Interestingly, *E. coli* laboratory strain MG1655, which lacks *crrA, crrB* and *crrC* genes, does not evolve resistance under these conditions (Fig. S4).

### Genetic basis of colistin resistance

We detected mutations in genes associated with colistin resistance in all experiments E1.1-E8.1 (Fig. 3). We did not detect any mutations associated with resistance in pre-colistin samples from pilot experiments P1.1 and P2.1 (Fig. S5). Thus, we surmise that almost all cells in the cultures tested were initially sensitive to colistin, and *de-novo* resistant variants were present in less than 1% of cells (detection limit of our WGS). We classified mutations as common (allele frequency *f* > 0.2) or rare (*f* < 0.2, i.e., present in less than 20% of sequenced alleles). Common mutations were observed in four genes: *pmrB, crrB, fimD*, and a hypothetical gene (BN373_30951) (AA changing mutation at 3242969, R630S). All these mutations, except *fimD* and BN373_30951, are known to increase resistance to colistin^8,24^. The mutation P43R in *fimD*, involved in the export and assembly of *fimA* fimbrial subunits across the outer membrane, were clonal (frequency *f* ≈ 1) at time point 3 in E1.1, E2.1, E5.1, E7.1 (4 out of 8 WGS samples) and subclonal at time point 2 in E5.1 and E7.1.

**Fig. 3.**
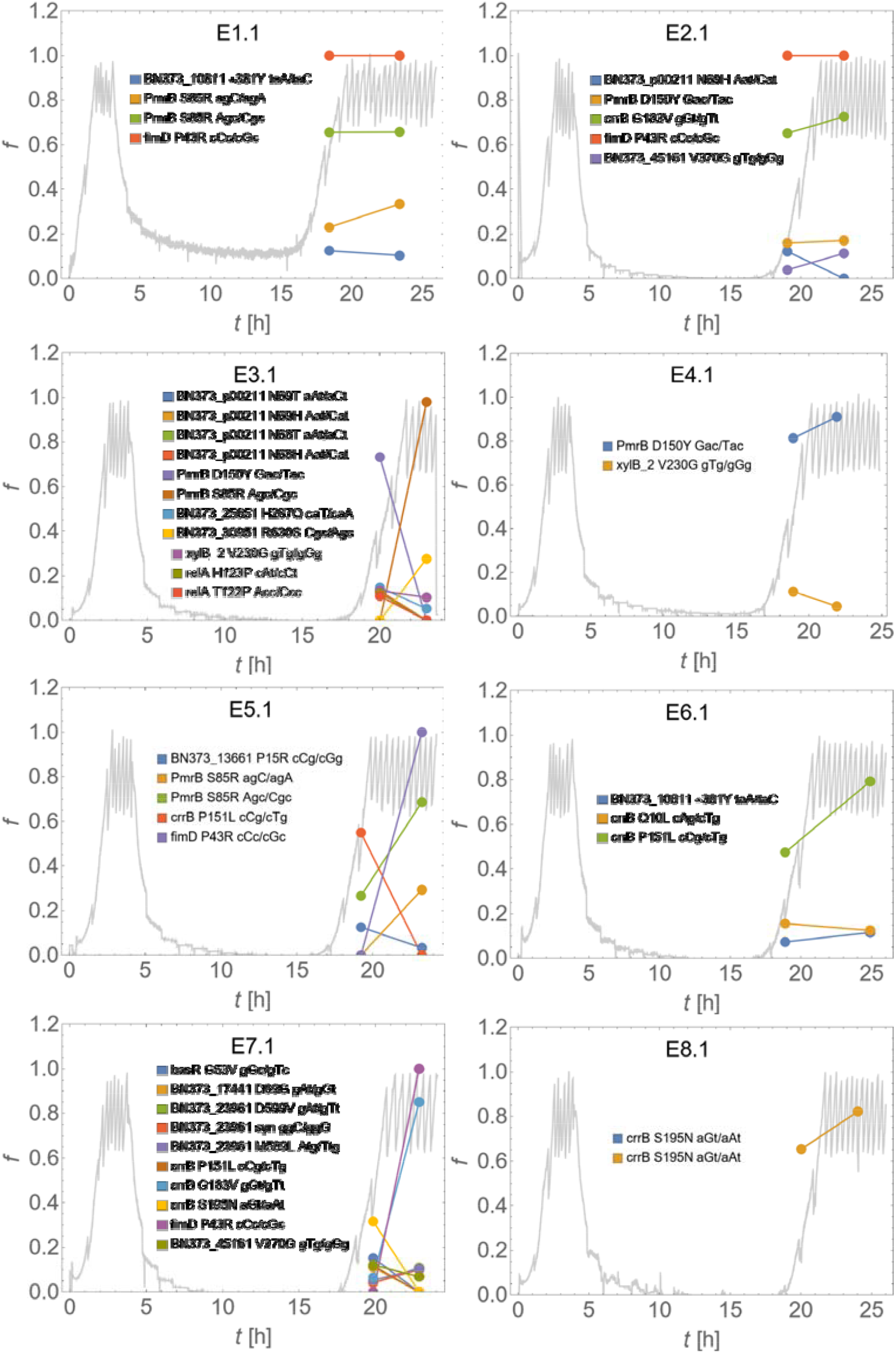
Fraction of alternative alleles (mutations) as a function of time, overlaid with the optical density curves. *f* is the allele frequency, *t* is time since the start of the experiment. Grey curve represents OD(t). Colours represent different mutations; their positions and gene names (if known, NCBI GenPept CDS locus tag if not) are listed next to each plot. Non-synonymous aminoacid substitutions have been also shown. “*” means a stop gained/stop lost type of mutation.

Rare mutations were found in genes not previously associated with resistance. We also found rare *pmrB, crrB, phoP, phoQ* variants; low frequency was likely caused by late occurence or lower fitness of these mutations.

Due to the limited number of WGS samples (n=8 + n=2 for the pilot experiments), some resistance mutations may not have been detected in our experiment. Based on the multiplicities of the observed putative resistant mutations, we estimated the number of possible but undetected mutations to be less than 12 (95% Bayesian credible interval), and most likely as little as two (Supp. Info.).

### Population dynamics of mutations and their relative fitness

We observed reduced genetic polymorphisms at time point 3 compared with time point 2 (Fig. 4), which suggested purifying selection acting against less-fit variants in the presence of colistin. For instance, mutations in *pmrB* (S85R, D150Y), occurred in 4 out of 8 sequenced experiments. However, *pmrB* D150Y occurred in E3.1 and E4.1 as the most frequent mutation at time point 2, but was replaced at time point 3 by *pmrB* S85R in E3.1. Thus, our data suggests that the mutation S85R in *pmrB* has a higher growth rate than D150Y in the presence of 10mg/L colistin. This was supported by bioinformatics analysis using PROVEAN^25^ (Supp. Info. “Fitness effects of mutations”).

**Fig. 4.**
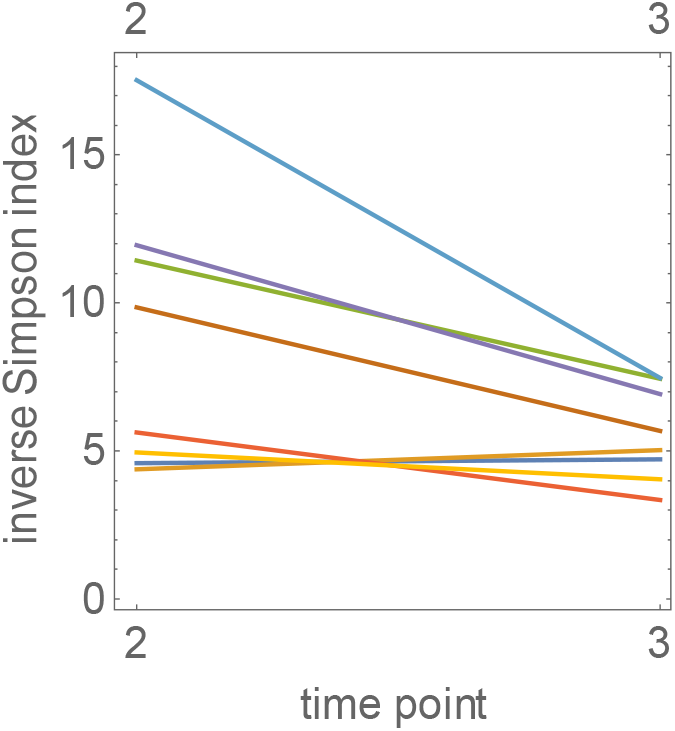
Genetic diversity decreases in time. Inverse Simpson index 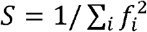 calculated using alternative allele (mutation) frequencies {*f*_*i*_} for time points 2 and 3. The numeric value of the index is approximately equal to the number of different alleles (mutations) in the population. The observed decay of 5 over time suggests selection acting on a set of alleles existing before time point 2.

Variations in *pmrB* (S85R), which arises as consequence of substitutions at either 253 A➔C or 255 C➔A, co-occurred with *fimD* P43R in E1.1 and E5.1. In both experiments, its frequency was close to f= 1, indicating that *pmrB* S85R and *fimD* P43R were present in the same clone. In E5.1, this combination replaced *crrB* P151L between time points 2 and 3. Therefore, we conclude that *pmrB* S85R + *fimD* P43R is more fit than *crrB* P151L at 10mg/L colistin. We observed three frequent (f> 0.2) mutations in *crrB* namely, G183V, P151L, S195N all of which have been reported in colistin resistant *K. pneumoniae*^6,8,24^. A combination of *crrB* G183V and *fimD* P43R outcompeted *crrB* S195N. Thus, we concluded that *crrB* G183V + *fimD* P43R is more fit than *crrB* S195N at 10mg/L colistin. Interestingly, *fimD* P43R always co-occurred with mutations in *pmrB* and *crrB*. In fact, PROVEAN analysis suggests that the P43R change in *fimD* is deleterious on its own (score −8.23, see the Supp. Info.). We hypothesise that this mutation increases the growth rate of resistant clones in the presence of colistin only in combination with *pmrB* and *crrB* mutations. In previous work, Cain et.al.^26^, report that *CrrB* can function as a regulator of fimbriae formation via *fimD*. However, our data show no differences in MICs in those clones, which harboured pmrB and *crrB* mutations either singly or in combination with *fimD* (Table 1) suggesting that the *fimD* mutation does not contribute directly to colistin resistance.

We also observed several mutations in *mgrB* (a missense variation, a stop gained, and a frameshift variation), albeit at very low frequencies: 2-5% at time point 2, 0-2% at time point 3 (E1.1-E8.1) and 4-6% (P1.1 and P2.1). Such a low prevalence of *mgrB* mutations agrees with the results of Cain et al^26^ and Janssen et al.^9^, who did not observe any *mgrB* mutants in liquid- and plate based colistin selection experiments^9,26^ of *K. pneumoniae*.

### Resistant mutants emerge before colistin treatment

Resistant cells, which cause regrowth in our experiments, could arise either before or during colistin challenge. In the case of pre-existing resistance, alternative allele frequency would have to be lower than ~0.01; otherwise WGS from time point 1 in experiments P1.1, P2.1 would have revealed the presence of such alleles in the population. To check if a small number of resistant mutants were present in the culture just before colistin exposure, we spot-plated 10µl of the culture onto LB agar with 10 mg/L of colistin. Each sample produced a few colonies, corresponding to ~ 10^4^ resistant cells per culture, suggesting that resistant variants exist in *K. pneumoniae* population before colisitin challenge.

### Mathematical model shows resistance is due to a small number of “founder cells”

We then used a mathematical model to verify our hypotheses about the origin of resistance (Methods). To obtain the unknown mutation rate, *µ*, we fitted the model to the observed distribution of regrowth times from E1.1-E8.4, and obtained *µ* = (0.6 … 1.2) x 10^−9^ (KS test p-value >0.01). Our model reproduces the experimental histogram accurately (Fig. 5). Dividing the estimated µ by the observed number of putative resistant mutations (2 in *pmrB*, 3 in *crrB*, 1 in *fimD*, and 1 in BN373_30951 gene) from WGS, we obtain an estimate of the per-base, per-replication mutation rate of *K. pneumoniae* as *γ*_*Kp*_ = (1 … 2) x 10^−10^. We are not aware of any direct measurement of *γ*_*Kp*_. However, *K. pneumoniae* mutates with approximately the same rate as E. coli in the fluctuation test^27,28^. Our estimate of *γ*_*Kp*_ is indeed similar to *γ*_*E*.*coli*_ ~2 × 10^−10^ from *E. coli* mutation accumulation experiments. Assuming that our estimate of *γ*_*Kp*_ is correct, the eight resistant mutations detected at the end point of the experiment must be a small fraction of all resistant mutations generated before colistin exposure. In particular, at least 26 point mutations in *mgrB* are known to cause resistance^30,31^. This should contribute at least ≈ 26x 1x 10^7−10^ ≈ 2.6 × 10^−9^ to the mutation rate *µ*, twice more than our upper estimate of *µ* based on the regrowth time. We postulate that *mgrB* mutations do occur in our experiment, but they are rapidly outcompeted by more fit variants. This does not require *mgrB* mutations to confer lower fitness in the absence of colistin^32^ but only to be less fit at 10 mg/L colistin; a similar behaviour has been reported for *E. coli* and fluoroquinolones^33^. Similarly, additional mutations in *pmrB, crrB*, and possibly other genes could theoretically occur but did not show up in WGS due to having a lower growth rate. This is supported by our observation that many more resistant cells are present in the culture just before colistin challenge, compared with ~ 10 − 50 “founder cells” predicted by the model (Fig. 6). Moreover, if we assume that there are six distinct resistant mutations, the model predicts on average *m*_>50%_ ~ 0.7 resistant mutations present in >50% of the cells, which is comparable to the actual number ~ 1 from the experiment (we exclude *fimD* and BN373_30951 since their role in colistin resistance is unclear). If the number of founder cells was much higher (either more distinct resistant mutations possible, or *γ*_*Kp*_ larger than assumed), *m*_>50%_ would be much lower (Fig. S6).

**Fig. 5.**
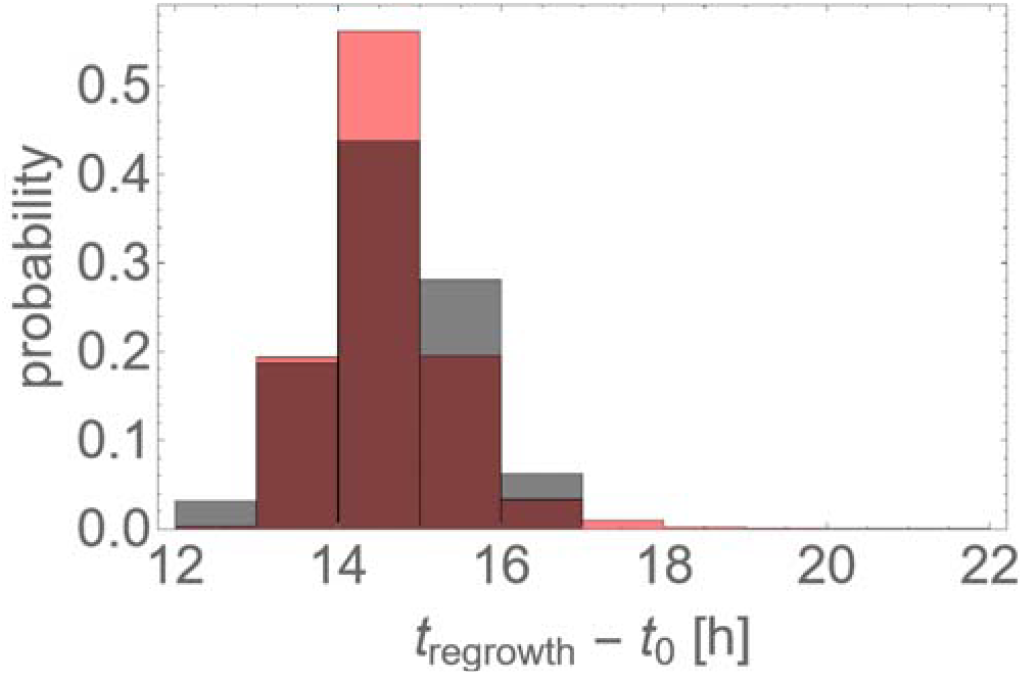
The model reproduces the histogram of regrowth times. Red = simulations with the best-fit mutation probability *µ* = 9 × 10^7−10^, gray = experimental data.

**Fig. 6.**
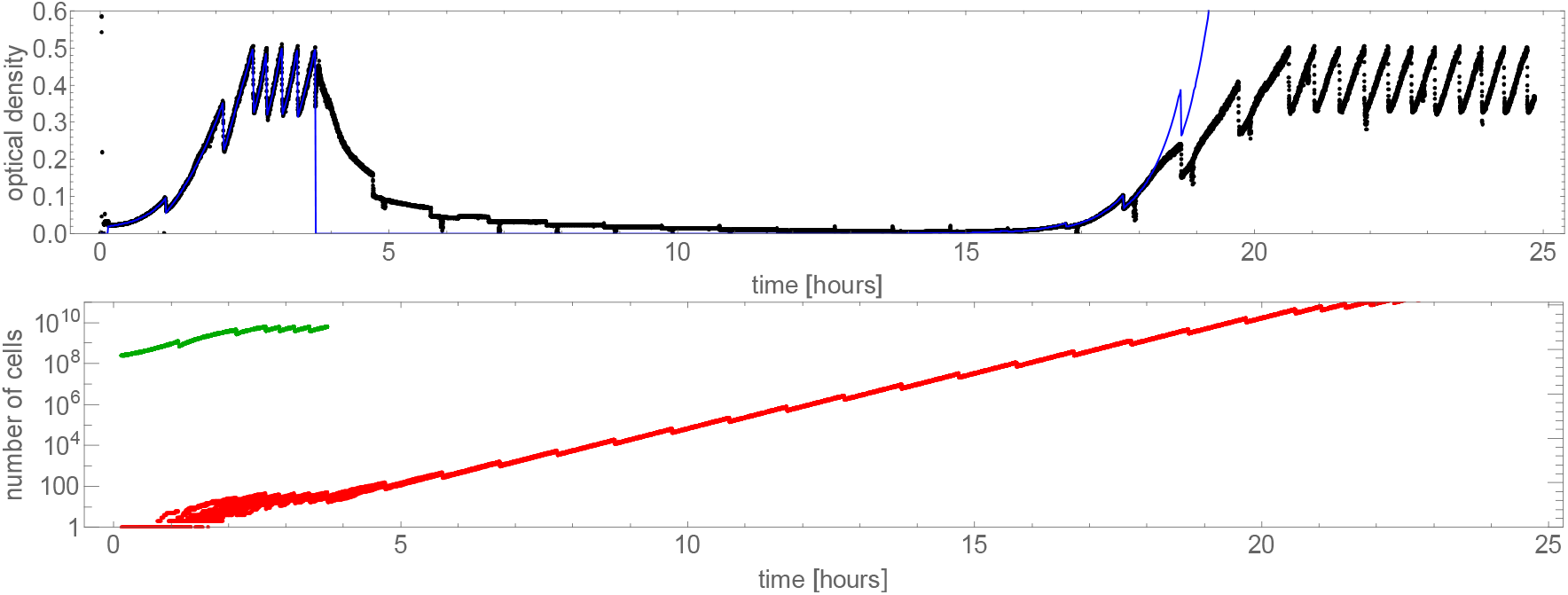
Dynamics of resistant mutants predicted by the computer model. Top: an example simulation run fitted to the OD versus time curve from the experiment E4.2 (black, blue = model). Bottom: number of live cells predicted by the model: green = colistin sensitive, red = colistin resistant. Different red curves correspond to different realisations of the computer simulation.

### Resistance evolution is likely in the clinical setting

We can now extrapolate the results from our bioreactor to a real infection. The only relevant parameters that affect the probability *P*_*resistance*_ in resistance evolution are the effective mutation rate *µ* at a clinically relevant concentration of colistin used here, and the number of divisions *N* of sensitive cells that have occurred before colistin treatment. Figure 7 shows *P*_*resistance*_ as a function of *N*, for the mutation rate *µ* = 0.9 × 10^−9^. The probability is close to zero for *N* < 10^7^, but approaches 100% for *N* > 10^9^. The number of divisions can be estimated from the bacterial load *L*, doubling time *t*_*D*_, and the duration of infection *T* before colistin is administered as 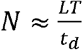. This assumes that all resistant mutants survive. If only a fraction of mutants survive, the probability of resistance will be proportionally smaller. We estimate (Supp. Info.) that *N* = 10^3^ − 10^14^, depending on the infection type, with numbers 10^5^ to 10^8^ for bacteraemia, and *N* > 10^8^ for lung infections^34^. Figure 7 shows that, at the high end of the bacterial load spectrum, resistance of *K. pneumoniae* to colistin is certain to evolve.

**Fig. 7.**
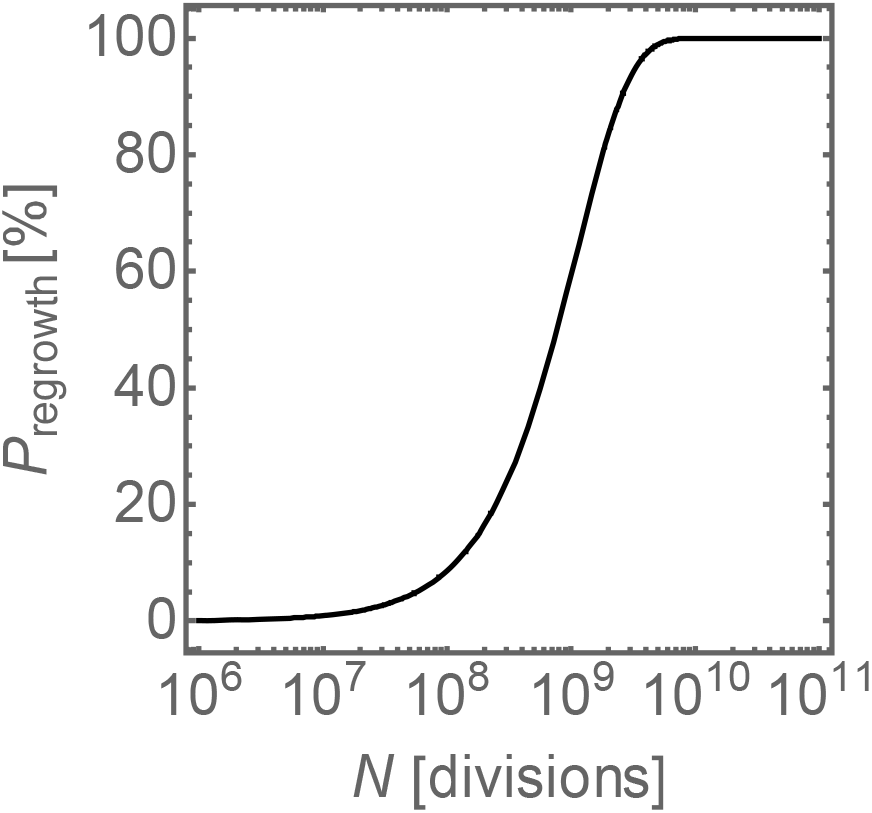
Model extrapolation to different bacterial densities. The probability of evolving *de-novo* resistance versus the number of divisions of *Klebsiella pneumoniae*, for *µ* = 0.9 × 10^−9^.

## Discussion

We used a bioreactor model of *K. pneumoniae* infection to better understand the origin and dynamics of resistance evolution to colistin. Although bioreactor models lack the complex host response of animal models, they benefit from higher reproducibility, better control and ability to monitor bacterial growth^35^. Compared to earlier work on AMR evolution in bioreactors^22,36^, agarose plates^26^, and batch cultures^9^, our approach (i) used clinically relevant combinations of pathogen and antibiotic concentrations, (ii) frequent bacterial density measurements, (iii) quantitative mathematical modelling, (iv) fully automated sampling at multiple time points, and (v) multiple replicates.

Our challenge experiments reveal rapid evolution of resistance, with MICs often exceeding 256 mg/L despite challenge at only 10 mg/L colistin. WGS shows that the evolved population is multiclonal but that in the presence of colistin the frequency of some alleles increase at the expense of other alleles, presumably due to fitness differences. Mathematical modelling reproduces the population dynamics of resistant mutants and shows that regrowth is derived from an existing population of resistant variants from an initially sensitive population. Less fit variants, including mutations in *mgrB*, are outcompeted by mutations in *crrB, pmrB*, which are more fit in these experimental conditions.

Consistent with the broader literature ^1,5,37^, our challenge experiments also demonstrate the emergence of variants harbouring mutations in both *crrB* and *pmrB* genes. Notably, in our experiments *mgrB* mutations occurred at an extremely low frequency. The lack of *mgrB* mutations is surprising but in line with other studies of plate or broth-based colistin challenge, which did not detect *mgrB* mutants^10,26^. In fact, recent data^10^ suggest that more than 30% of colistin-resistant clinical isolates have mutations in *crrAB* rather than in *mgrB* locus.

Given that the 10mg/L colistin used reflects a humanised dose^4,11,12,35^, and considering that the ratio between therapeutic concentrations and the MIC is on the order of 5-10 (as much larger concentrations are toxic to the patient)^15^, our study suggests that *de novo* mutations which confer colistin resistance are likely to occur in the population of *K. pneumoniae* during treatment. Importantly, our data suggests that a single, spontaneously generated mutation can increase the MIC 100x or more in a non-strain dependent manner supporting the rationale against using colistin and other antimicrobial peptides as a monotherapy for *K. pneumoniae* infections.

## Supporting information

Supplementary Table - coding SNPs

## Acknowledgements and Funding

TS acknowledges financial support from the Medical Research Council (MR/P007597/1) and BW acknowledges the support of the project financed under Dioscuri, a programme initiated by the Max Planck Society, jointly managed with the National Science Centre in Poland, and mutually funded by Polish Ministry of Science and Higher Education and German Federal Ministry of Education and Research (grant no. UMO-2019/02/H/NZ6/00003).

## Supplementary figures

**Fig S1.**
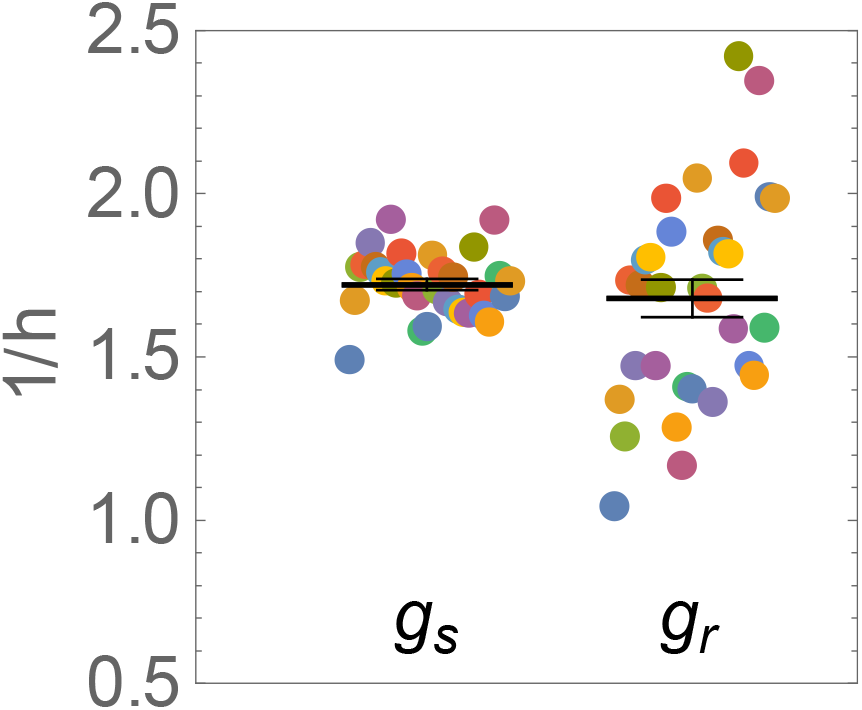
Growth rates of sensitive (*g*_*S*_, average before colistin) and resistant (*g*_*R*_, max. growth rate from the regrowth phase) populations. Different colours represent different experiments (32 in total). Horizontal lines are mean +/- S.E.M.

**Fig. S2.**
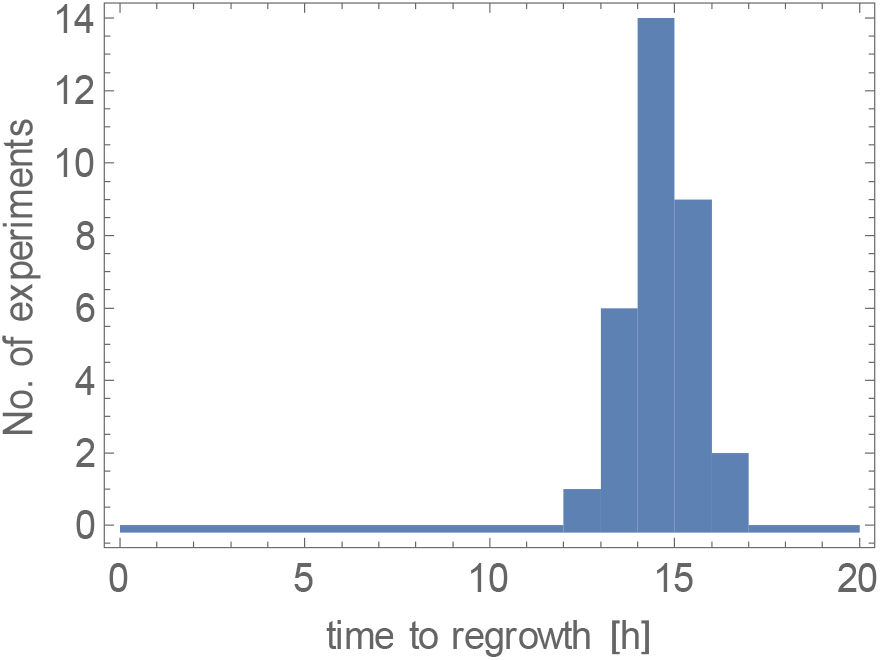
Distribution of regrowth times (bin size = 1h). The time to regrowth is defined as the time from the first injection of colistin until the time at which OD increases above 0.1.

**Fig. S3.**
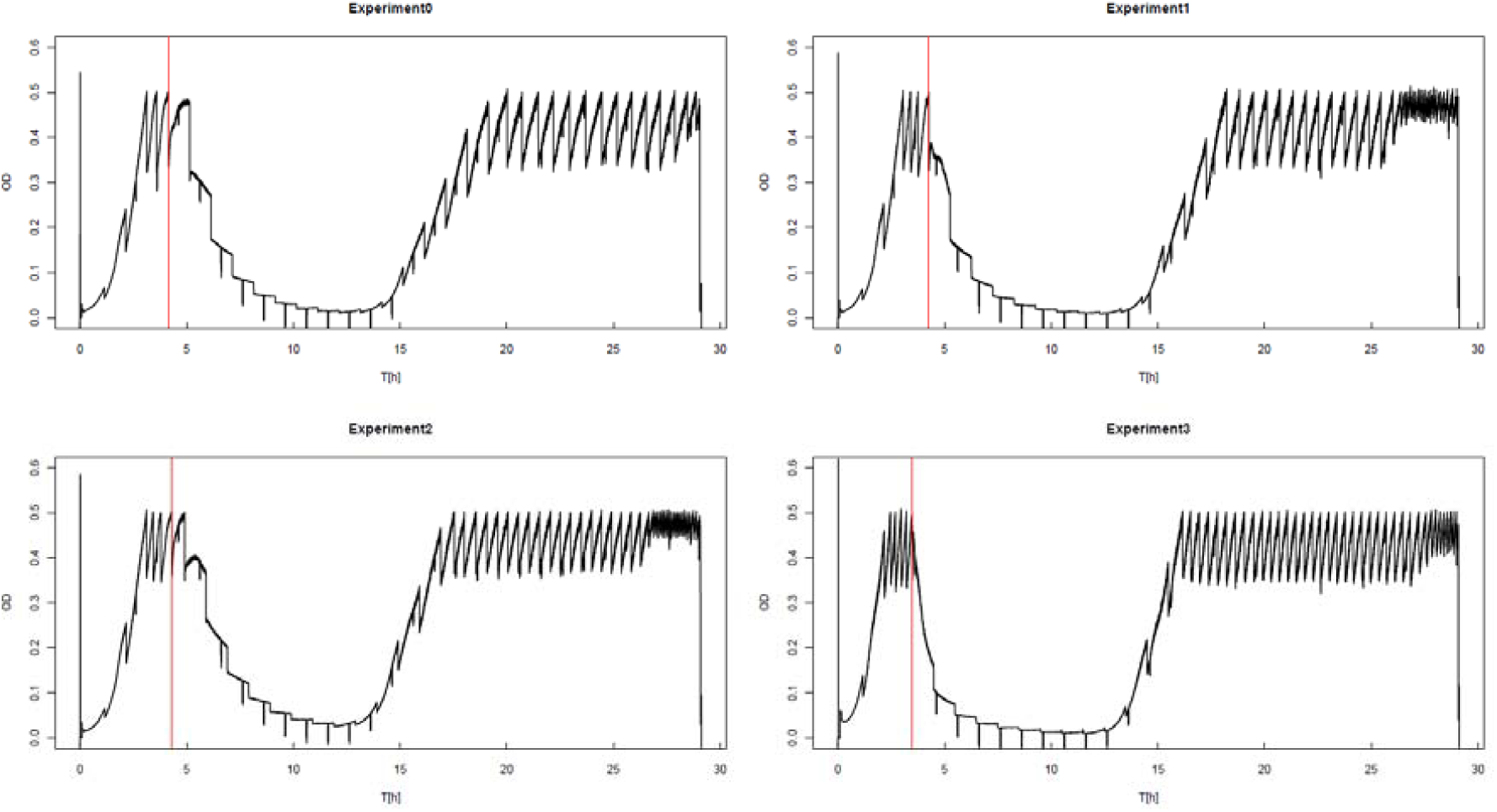
OD vs time for *K. pneumoniae* 52145, a K2:O1 (hypervirulent) strain. Red line indicates the beginning of colistin treatment.

**Fig. S4.**
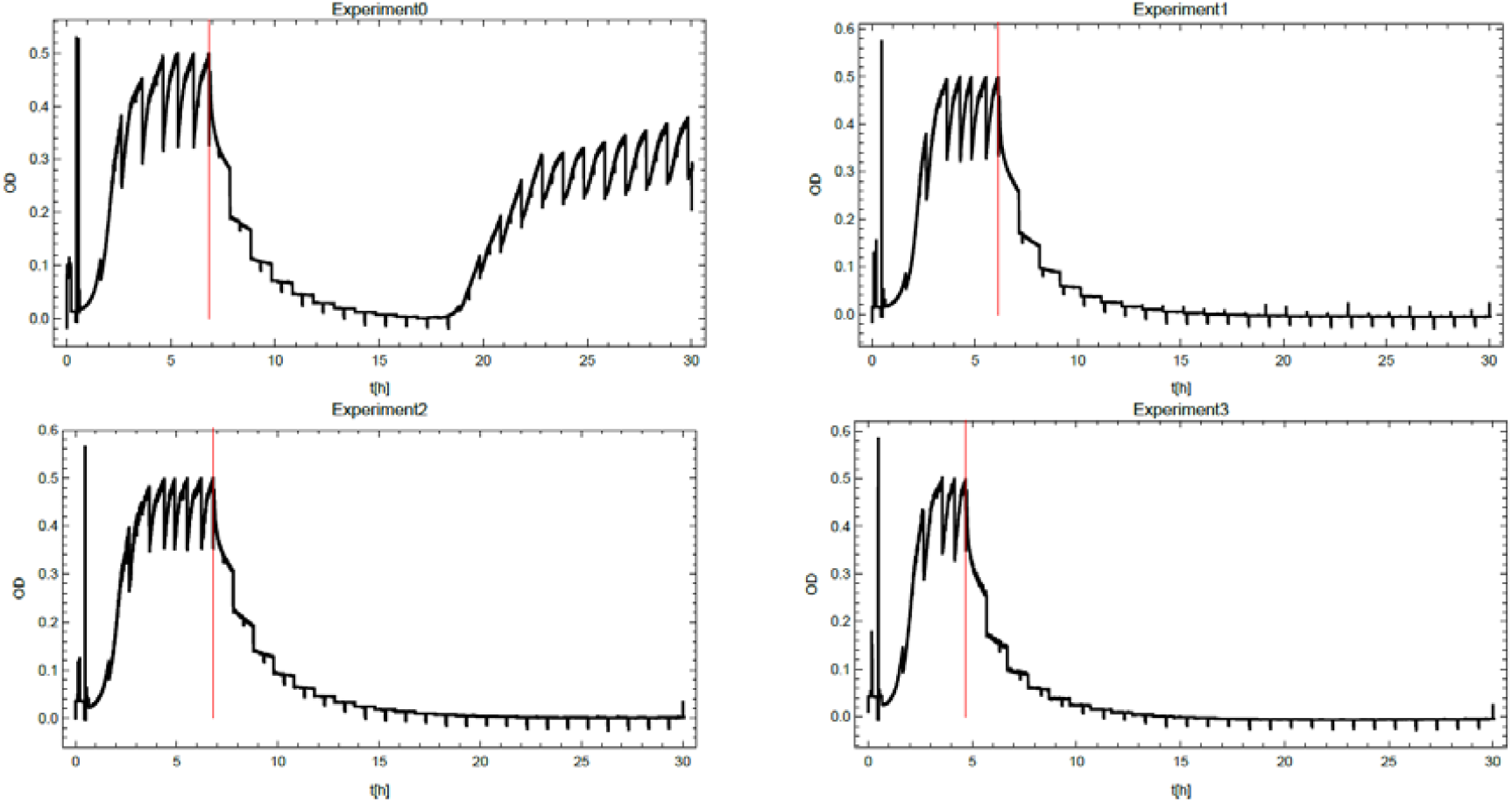
OD vs time for four experiments: Ecl8 (panel 1) and *E*.*coli* MG1655 (panels 2-4). Red line indicates the beginning of colistin treatment.

**Fig. S5.**
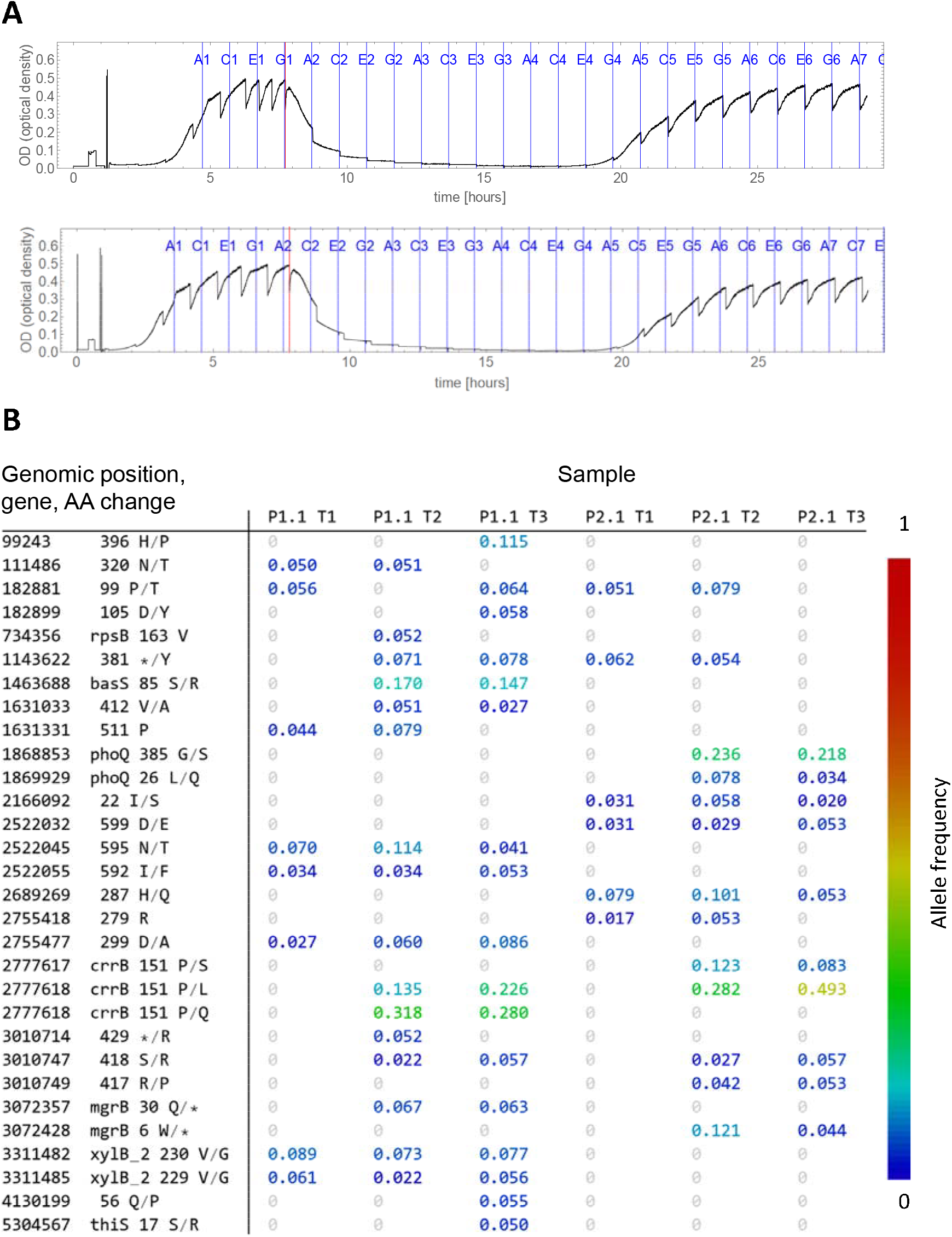
(A) OD vs time for two out of eight experiments from the pilot runs P1 and P2. Red line indicates the beginning of AB treatment. Blue lines with well numbers show the times at which the culture was sampled by the autosampler. We performed WGS for E1, E5, A7 (P1) and E1, E5, A7 (P2). (B) Frequency of alternative alleles (mutations) in all samples. Pre-colistin samples show only low-frequency (<10%) mutations in genes not related to colistin resistance.

**Fig. S6.**
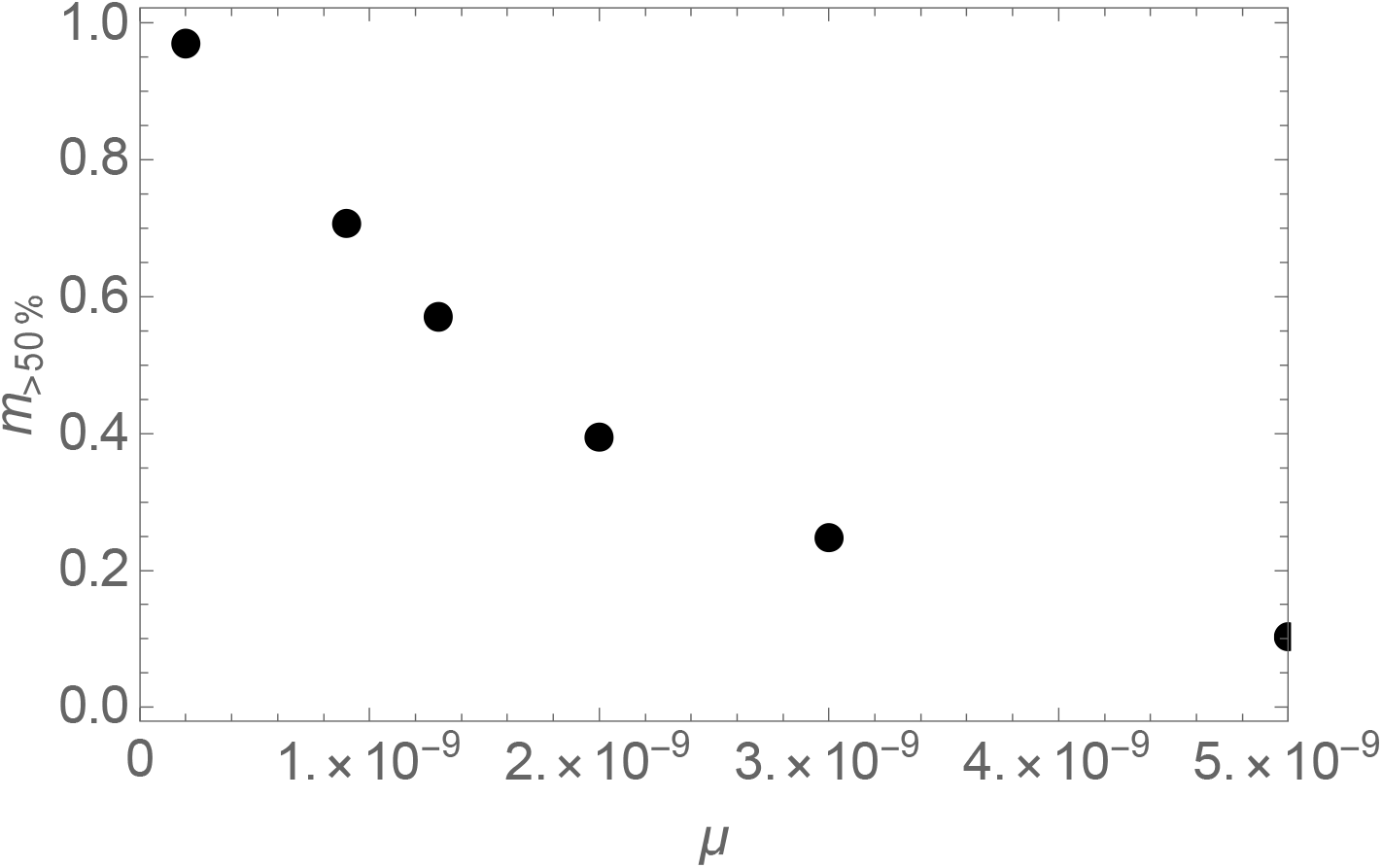
The number of resistant mutations present in >50% of the cells at the end of the experiment, as predicted by the model, for different mutation rates *µ*. The actual number determined from 8 experiments is 1 mutation/experiment.

