## Supplementary Information for "Rapid Evolution of Colistin Resistance in a Bioreactor Model of Infection of *Klebsiella pneumoniae*"

**More details about the bioreactor**

Bioreactor *in vitro* models of infections[1–3] have several advantages over animal models: (i) better control over experimental conditions, (ii) bacterial growth can be monitored in time resolved manner, (iii) samples can be collected non-destructively, stored and analysed at any given specific time. The main disadvantage is the absence of the complex host response to the infection. However, as evolution of resistance to antibiotics, bacterial load and the mutation rate are the primary factors, the lack of the immune system can be then compensated to some extent by lowering the density of bacteria.

Our approach enabled us to i) track the bacterial growth in a time-resolved manner before and after exerting colistin pressure and ii) to characterize phenotypic and genetic (via whole-genome sequencing, WGS) response to colistin at different time points of the regimen.

Compared to earlier work on related *in vitro* models[4–7],[8] our approach enables (i) full automation, thus experiments can be run without interruption for >24h, (ii) high-frequency OD measurements, (iii) four biological replicates run in parallel, (iv) quantitative comparison with mathematical modelling thanks to a relatively simple dilution protocol that can be easily accounted for in the mathematical model.

**OD calibration**. Before each experiment we calibrated the OD sensors using two point calibration (low $OD_{600}$=0.1 and high $OD_{600}$=0.5) by inserting two bottles (LB and a *K. pneumoniae* Ecl8 culture of known $OD_{600}$) into each holder, and executing an appropriate calibration procedure in the software. The software assumes a linear relationship between the current from a photo-transistor measuring the amount of scattered light, and the true $OD_{600}$. We tested linearity by measuring the signal for colloidal suspensions of known $OD_{600}$ (up to $OD_{600}$=0.5). While the actual relationship is not perfectly linear (in particular, for $OD_{600}$>0.3), the deviation from linearity is less than 10% for the whole range of OD values used in the experiment.

**Antibiotic (AB) concentration control**. The software calculated the amount of time the LB and AB pumps should run for to mix both liquids, and the liquid present in the bottle, in proportions required to achieve a desired concentration of colistin. Since the software controlled the running time of the pumps and not the actual volume added, the pumps were calibrated before each experiment by measuring the time it took to deliver a known volume.

**Preparing the system.** Media reservoirs, culture bottles, plastic tubes were autoclaved and aseptically assembled to avoid contamination. LB (Formedia) was prepared and autoclaved prior to filling the 2L LB reservoir bottle, and a smaller 1L antibiotic reservoir bottle. Colistin was added to the AB reservoir bottle to appropriate concentration (50 µg/mL). After assembling the system, tubes were primed with LB and AB by activating the pumps in the control software. Each bottle was then filled with LB which was subsequently drained out by manually opening the tube and carefully pouring out the liquid, and the procedure was repeated several times to get rid of AB injected into the bottle during priming. The incubator was switched on, and the system left for at least 1h for the temperature of all components to stabilise at 37 °C. After the experiment, the system was cleaned by replacing the reservoir bottle with tap water, then PRESEPT (2,500 ppm sodium dichloroisocyanurate) as per manufacturer's instructions, then tap water again, and pumping it through the tubes and culture bottles, and finally disassembled and autoclaved.

**Temperature and OD in the bioreactor.** The temperature remained within the 36-37℃ range for the duration of all experiments. The average $OD_{600}$ (0.35-0.5 in the pre-treatment phase) was higher than in other evolutionary experiments[2,3,9] (OD600=0.1 to 0.2). However, we decided to use higher cell density in our experiments for several reasons, firstly, it provides a larger pool of initial mutations and secondly, growth is slower at higher cell densities, which reflects the stringent physiological conditions which bacteria potentially encounter *in vivo*.

### WGS/Data Analysis

**DNA isolation**. From each sample point, we extracted genomic DNA using Lucigen Kit (Bioresearch, Lucigen. UK). Briefly, each sample was collected from the 96-well plate and centrifuged at 13, 000 rpm X 10 min. Supernatant was discarded and resuspended in 300 µL of Lysis buffer supplemented with 1 µL of proteinase K (20mg/mL) and incubate for 15 minutes at 65 ℃ with shaking at 14, 000 rpm. Samples were then cool down to 37℃ and 2 µL of RNAse (10mg/mL) was added followed by 30 min incubation at 37℃. Afterwards, samples were incubated on ice for 5 min and 175 µL of protein precipitation buffer was added. Samples were vortex for 10s and spun down at 13, 000 rpm for 10 min. Supernatant was transferred to a clean tube and 500 µL of 100% isopropanol were added to precipitate genomic DNA. Supernatant was discarded and washed twice with 500 µL of 70% ethanol. Finally, we discarded supernatants and allowed pellets to air-dried before resuspending in 50 µL of molecular biology grade water. We quantified and checked for DNA integrity using a TapeStation (Agilent, UK) according to manufacturer's instructions. All samples with DNA Integrity Number (DIN)≥8.0 were used for WGS analysis.

**WGS data analysis.** Samples were aligned to the *K. pneumoniae* Ecl8 reference genome and variants called with the software package Snippy v4.3.6 (https://github.com/tseemann/snippy) with minimum depth for a base call of 3X, minimum alternate allele frequency 0.01, and Freebayes parameter '--pooled-continuous'. Multi-allelic variants were decomposed and alleles normalized with the vt package v0.5 (http://bioinformatics.oxfordjournals.org/content/31/13/2202). Reference allele, alternate allele, and total depth information was extracted for each variant site. Quality control metrics for reads and alignments were calculated with FastQC v0.11.4 (https://www.bioinformatics.babraham.ac.uk/projects/fastqc/) and samtools v1.6 flagstat (https://www.ncbi.nlm.nih.gov/pubmed/19505943). Depth at every base in each sample across the reference genome was calculated with bedtools v2.21.0 genomecov. We calculated the fraction of alternative alleles in time points 1, 2 and 3 using OVC as a reference. Single nucleotide variant sites where the alternate allele frequency was at least 0.05 greater than the OVC sample for the experiment were selected. A region around 142Kbp (142203-142448) was excluded due to low coverage and quality.

**Estimating the number of putative resistant mutations.** We observed the following mutations in *pmrB*, *crrB*, *fimD*, and *BN373_30951*, and their multiplicities (in how many replicate experiments a given mutation occurred):

| Mutation | *pmrB* S85R | *pmrB* S85R | *fimD* P43R | *crrB* G183V | *crrB* P151L | *crrB* S195N | *pmrB* D150Y | *BN373_30951* R630S |
|---|---|---|---|---|---|---|---|---|
| Multiplicity | 2 | 3 | 4 | 2 | 1 | 1 | 1 | 1 |

It is possible that more distinct resistant mutations could have been generated in our experiment but we did not observe them because of the limited number of sequenced samples (*n*=8). To estimate how many mutations we could have missed, we used Bayesian inference. We simulated 1,000,000 realizations of our experiment, assuming that $M$ distinct resistant mutations could occur, each with equal probability. In each simulation, we created a vector of multiplicities $\{n_1, n_2, \dots, n_m\}$ by randomly choosing $N$ out of $M$ mutations and counting the number of occurrences of each mutation. If the length $m$ of this vector was the same as the length $n = 8$ of the experimental multiplicity vector, and the two vectors were identical, we considered this simulation to be a "success" – the simulated multiplicities were consistent with the experimental multiplicities. We then counted the number of successes in one million trials. Repeating this procedure for different $M$ we obtained the following plot of the number of successes versus $M$, which is (modulo a normalization factor) the Bayesian posterior probability of the number of putative mutations:

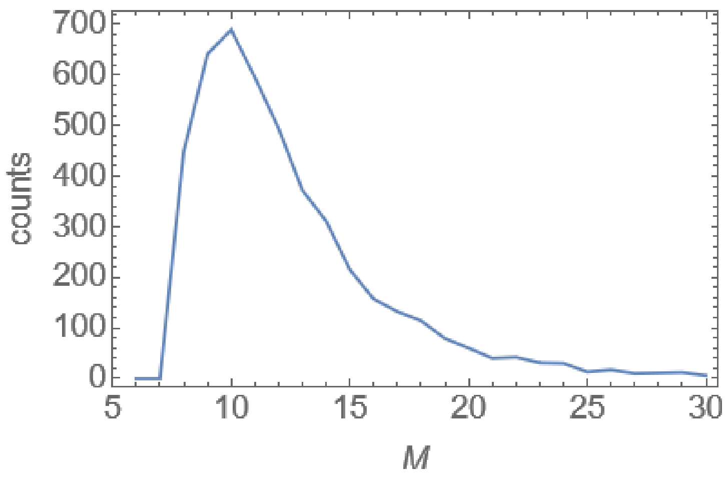


Using the posterior probability we estimated the 95% credible interval for $M$ to be between 8 and 20. Since we observed 8 mutations, the upper limit on the number of missed mutations is 12 with probability 95%. However, the most probable $M = 10$, which means we have most likely missed only two mutations.

**Fitness effects of mutations.** PROVEAN PROTEIN online software (http://provean.jcvi.org/) was used to computationally predict the fitness of point mutations found in WGS. The software calculates a score based on the reference and variant versions of amino-acid sequences by comparing them with sequence homologs from the NCBI protein database. We used the following versions of the FimD and PmrB proteins (blue symbols denote amino-acids that have been altered in the mutated variants):

Using the prediction cut-off of -2.5 as suggested by the software authors, we obtained the following scores:

| Protein | Variant | PROVEAN score | Prediction |
|---|---|---|---|
| FimD | P43R | -8.24 | deleterious |
| PmrB | S85R | -2.96 | deleterious |
| PmrB | D150Y | -8.48 | deleterious |

The results for the mutations that we found were in accordance with those seen in the literature[10–12]. The results also agree with phenotypic effect observed (Main text, Table 1).

**Model parameters for real infections.**

All parameters in the formula for $N$, the number of bacterial divisions, are difficult to estimate *in vivo*. However, $t_D$ and $T$ cannot vary by more than a small factor due the biology of *K. pneumoniae* and the limited duration of a typical infection: $t_D$ cannot be smaller than 20 mins and longer than a few hours, whereas $T$ is likely to be between 10h and a few days. On the other hand, $L$ can potentially vary by orders of magnitude: a few thousand cells may be responsible for bacteraemia, whereas $L$ can be easily over $10^7$ in urinary infections, and potentially more than $10^{11}$ in lung infections, see Table S2 below. Assuming $t_d = 0.3 \dots 2$ h, $T = 10 \dots 100$ h, and using the formula $N \approx \frac{LT}{t_d}$, we get a very large range of $N = 10^3 - 10^{14}$. For bacteraemia, assuming $L = 10^3 \dots 10^7$, we get $N = 10^5$ to $10^8$. For lung infections, the numbers can be much higher since $L > 10^7$ is typical.

| **Infection** | **Bacterial density (cells per sample volume)** | **Volume of infected tissue (order of magnitude)** | **Total bacterial load $L$ (cells)** |
|---|---|---|---|
| Lung, mountain gorilla[13] | $10^7$ per microliter of mucus | 100 ml | $10^9$ |
| Lung, human[14] | $10^4$ to $> 10^7$ per ml of sputum | 100 ml | $10^6 - 10^9$ |
| Elderly person, urine[15] | $> 10^5$ per ml of urine | 100 ml | $> 10^7$ |
| Urine, human[16] | $10^2$ to over $10^5$ per ml | 100 ml | $> 10^4$ |
| Bacteremia, adults, blood[17,18] [ref] | $1 - 10^3$ per ml blood | 5000 ml | $10^3 - 10^7$ |
| Mice, infection model, blood[19] | $10^6$ per ml blood | 2 ml | $10^6$ |
| Mice, infection model, lung | $10^5 - 10^6$ per ml mucus[16,19]<br>$10^{11}$ per ml[14,20] | 1 ml | $10^5 - 10^{11}$ |

**Table S2. Estimation of the bacterial load $L$ for different infections.**
